# FLIM Imaging of mCherryTYG Deciphers pH Dynamics and Lifestyles of *Salmonella* Typhimurium

**DOI:** 10.1101/2025.01.27.635066

**Authors:** Moirangthem Kiran Singh, Marion Fernandez, Rahul Dilawari, Parisa Zangoui, Linda J. Kenney

## Abstract

Intracellular pH regulation is fundamental to bacterial adaptation, virulence, and survival in diverse environments. *Salmonella* Typhimurium, a key human pathogen, exploits host and environmental pH cues to transition between planktonic, biofilm, and virulence-associated states. However, precise tools to monitor bacterial pH dynamics at subcellular resolution have been limited. Herein, we report the development and application of mCherryTYG, a genetically encoded pH-sensitive fluorophore optimized for fluorescence lifetime imaging (FLIM), enabling robust and high-resolution pH measurements across diverse conditions. mCherryTYG demonstrated exceptional sensitivity across a broad pH range (5.5–8.5) with consistent lifetime responses, and was unaffected by temperature, buffer composition, or ionic strength. Using FLIM, we characterized the pH dynamics of *Salmonella* across *in vitro*, host, and biofilm contexts. Under acidic stress *in vitro*, *Salmonella* maintained a uniform intracellular pH (∼6.04), providing clarity on previously debated heterogeneity. In infections of HeLa cells, *Salmonella* existed in distinct pH environments: acidic vacuolar pH (∼5.89) and neutral cytoplasmic pH (∼7.10). During the late infection stage, ∼17% of the bacterial population retained an acidic pH. Biofilm studies revealed stratified pH profiles, with acidic pH near the bottom and neutral pH at the surface, mirroring patterns observed in other pathogens. In heterologous host models, pH gradients shaped bacterial adaptation strategies. In *C. elegans*, *Salmonella* experienced a progressive pH gradient from neutral pH (∼7.10) in the anterior lumen to acidic pH (∼6.45) in the posterior. Similarly, in zebrafish, *Salmonella* encountered acidic lysosome-rich enterocytes (∼5.84) and neutral regions (∼7.33) in the anterior gut. This study establishes mCherryTYG-FLIM as a transformative tool for studying bacterial pH regulation, revealing pH as a critical modulator of *Salmonella* lifestyle transitions: virulence, and persistence. Our findings provide new insights into host-microbe interactions and present pH as a promising target for therapeutic interventions against bacterial infections.

## INTRODUCTION

The pH level within cells plays a crucial role in their biological functions and overall metabolism. Inside eukaryotic cells, intracellular pH plays a fundamental role in cell functions such as cell growth and metabolism ^[1]^, ion transport and homeostasis ^[2]^, endocytosis^[3]^, post-translational protein modifications^[4]^ and signaling cascades^[5]^. Abnormal shifts in pH levels are also evident in cellular apoptosis and diseases such as cancer, as well as neurological conditions such as stroke and ischemia, among others.^[4, 6, 7]^ In bacteria, pH impacts the growth, nutrient uptake, and ability to thrive in different environments. Unlike eukaryotes, Gram-negative bacteria demonstrate the capacity to endure considerable fluctuations in osmolality (up to 1800LmOsmLKg^-1^), withstand substantial decreases in cytoplasmic pH (to 5.5 or lower), and exhibit resilience in diverse environments both within and outside the host.^[8, 9]^ Pathogenic bacteria often exploit specific pH environments to colonize the host and evade its immune system. A notable example is *Salmonella enterica* serovar Typhimurium, which has evolved survival tactics to flourish within membrane-bound acidic compartments of macrophages, or to form biofilms, exploiting acid pH as an important environmental signal.^[10–12]^

Given the pivotal role that pH sensing plays in bacterial pathogenesis and lifestyle adaptations, the development and enhancement of pH probes represents a dynamic area of research. Numerous organic fluorophores sensitive to pH, such as BCECF^[10,18,21]^, SNARF ^[21,22]^, or fluorescein ^[22]^, have been developed and employed extensively for spatially resolving intracellular pH and dynamics in biological imaging studies. However, despite their extensive application, their effectiveness is hindered by several challenges, including dye loading, leakage, and inadvertent binding to proteins and membranes. One significant challenge associated with organic dye-based pH sensors is accurately controlling their localization within the cell, as well as managing potential toxicity and interference with cellular metabolic activities, thus limiting their utility.^[13, 14]^ An alternative strategy involves employing non-toxic, genetically-encoded fluorescent biosensors such as pHluorin^[15]^ and SypHer^[16]^, which enable precise targeting to specific subcellular locations. Other informative fluorescent biosensors include Cy11.5 and pHlameleons^[17]^, mCherryEA^[18]^, pHTomato^[19]^, pHuji^[20]^, mNectarine^[21]^, among others. These biosensors function on a ratiometric principle, where the pH at each pixel is calculated based on the ratio of fluorescence intensities measured at two emission wavelengths after excitation at two specific wavelengths. However, a significant limitation of ratiometric pH sensors is the decreased signal-to-noise ratio at low fluorescence intensities. This issue arises from the intrinsic error amplification associated with ratio calculations and the varying focal depths of microscopy at different wavelengths. Furthermore, this method relies on excitation power and filter bandwidths; separating pH measurements from the sensor concentration presents another challenge. Many of these limitations can be addressed by utilizing fluorescence lifetime imaging microscopy (FLIM) of lifetime-based pH sensors. Fluorescence lifetime is the average duration a fluorescent molecule spends in its excited state before emitting a photon and returning to its ground state. It represents an intrinsic property (in the absence of non-radiative processes) of the fluorophore, offering insight into the environmental factors influencing its photophysical properties. Given its sensitivity to the microenvironment, changes such as fluctuations in pH can be accurately assessed using FLIM. FLIM serves as an excellent alternative to ratiometric measurements, because the life-time is independent of fluorophore concentration and excitation power, requiring only a single excitation wavelength/emission interval, and enabling accurate pH evaluation without the need for a second reference fluorophore.^[22]^

In this study, we utilized a FLIM-based approach to address challenges associated with dual excitation and leverage the fluorescence lifetime properties of mCherryTYG, a variant of mCherry, as a FLIM-based pH sensor.^[23]^ Wildtype mCherry is frequently employed as a red fluorescent protein (RFP) due to its minimal cytotoxicity. This characteristic makes it an excellent foundation for the development of an RFP pH sensor. The fluorescence of wildtype GFP, a similar protein, is known to be stable from pH 6-10; it is plausible that mCherry would behave similarly. The development of mCherry variants with pH sensitivity could potentially allow the monitoring of intracellular pH changes. Using time-resolved fluorescence spectroscopy (TRFS), an mCherryTYG variant (M66T), exhibited a remarkable 16-fold increase in brightness when the pH was shifted from 5.5 to 9.0, displaying notable pH sensitivity.^[35]^ In contrast to TRFS, FLIM offers spatial resolution in addition to temporal fluorescence lifetime data, allowing the visualization of lifetimes across different regions of a sample. This enables the detection of heterogeneities in complex environments, which TRFS lacks, as it provides only bulk measurements. Herein, we establish that mCherryTYG serves as an effective FLIM-based RFP pH sensor. It demonstrated insensitivity to laser intensity or protein concentration, while exhibiting sensitivity to pH changes with both temporal and spatial precision, thereby enabling quantitative pH measurements in live cell imaging. We extensively characterized the pH-dependent lifetime properties of the mCherryTYG mutant protein, both in solution and within live bacterial cells. To illustrate its utility in understanding bacterial pathogenesis and adaptive responses, we mapped and generated detailed spatiotemporal profiles of pH fluctuations within bacterial cells exposed to varying pH conditions. This mapping encompassed a diverse array of environments, including *in vitro* settings and *in vivo* scenarios such as *Salmonella*-containing vacuoles during the course of bacterial infection, in biofilms, and the heterologous host environments of nematodes and zebrafish. Our findings underscore the critical role of bacterial pH regulation in both bacterial pathogenesis and in the adaptation of bacterial lifestyles.

## EXPERIMENTAL SECTION

### Materials

Unless specified otherwise, all experiments were performed using molecular biology grade chemicals. Cell culture media and supplements were sourced from Thermofisher (Invitrogen). Chemicals and oligonucleotides acquired from commercial suppliers were used without additional purification.

### Strains, plasmids, and molecular biology

*Salmonella enterica* serovar Typhimurium strain 14028s is the wild-type strain, and various isogenic mutants are described in the text. The plasmid pFPV-mCherry, generously provided by Olivia Steele-Mortimer of Rocky Mountain Labs, Hamilton, MT (Addgene plasmid #20956), was used. Additionally, pRSET-B mCherry was a gift from Kalina Hristova (Addgene plasmid #108857). Site-directed mutagenesis was used to modify the wildtype mCherry gene, resulting in mCherryTYG (M66T), where the chromophore methionine was substituted with threonine. The bacteria were cultured over-night in LB broth at 37°C with agitation at 250 rpm.

### Protein expression and purification

pRSETb-mCherryTYG was introduced into BL21(DE3) cells, cultured in 250 mL of LB me-dia at 37°C for 20 h and stored at -20°C. The protein was isolated using HisTrap Nickel col-umns (GE Healthcare) following the manufacturer’s protocol. Subsequently, the purified protein was dialyzed using a BioSpin30 column (BioRad) into a storage solution consisting of 5 mM 3-(N-morpholino)propanesulfonic acid (MOPS), 150 mM NaCl, and 5% glycerol at pH 7.4, and then stored at 4°C. A single amino acid substitution at mCherryTYG (M66T) was further confirmed using mass spectrometry (see Figure S1).

### Characterization of mCheryTYG in calibration buffer

For characterization of purified mCheryTYG, the protein was dialyzed using the BioSpin30 into the corresponding calibration buffers. Protein concentration was measured using a Nanodrop (DeNovix DS-11). Buffers with specific pH values for calibration using purified proteins were prepared according to the previously established protocol.^[23]^ Buffers contained 150 mM 2-(N-morpholino)ethanesulfonic acid (MES), pH 5.5 and 6.0; 50 mM MOPS, 50 mM Tris, and 50 mM Bis-Tris, pH 6.5, 7.0. 7.5, 8.0 and 8.5. Each buffer solution was adjusted to the desired final pH using 1 M NaOH or 1 M HCl at 37°C.

### pH calibration in live bacterial cells using clamping buffers

To convert fluorescence life-time into pH, a calibration curve was established for the pH range of 5.5 to 8.5 using live bacterial cells expressing mCherryTYG. Briefly, *Salmonella* Typhimurium strain harboring pFPV::mcherryTYG was cultured overnight in 1 mL of LB media, followed by washing with MgM and resuspension in 50 μL of MGM at pH 7.4. This solution was then used to inoculate MgM at pH 7.4 at a dilution of 0.1%, and the bacteria were incubated at 37°C for 2 h. For clamping, the bacteria were resuspended in clamping buffer at various pH levels (composed of 0.4 g/L KH2PO4, 2 g/L (NH4)2SO4, 7.45 g/L KCl, and 50 mM MES or Tris Base or HEPES) supplemented with 40 μM nigericin (Sigma Aldrich) and incubated for 1 h at 30°C.^[9, 24]^ For imaging, bacteria were seeded on an IBIDI chamber slide coated with poly-L-lysine.

### Measurement of mCheryTYG pH in *Salmonella*

Bacteria from an overnight LB culture were inoculated at a 1:100 ratio into LB at pH 4.5, MgM at pH 5.6, and MgM at pH 7.2. The cultures were incubated until the optical density at 600 nm (OD_600_) reached approximately 1. We then harvested *Salmonella* cells and seeded them onto Poly-L-Lysine coated 18-well chambered slides. The slides were centrifuged at 2500 rpm for 10 minutes at room temperature, washed, and then filled with the corresponding conditioned media.

### Mammalian cell culture

The human epithelial cell line HeLa (ATCC CCL-2, obtained from the American Type Culture Collection) was used for *Salmonella* infection experiments. HeLa cells were maintained in high-glucose Dulbecco’s Modified Eagle Medium (DMEM; 4.5 g/l glucose) supplemented with glutamine, pyruvate (Thermo Fisher), 10% heat-inactivated fetal bovine serum (FBS; Thermo Fisher), and a 1× concentration of antibiotic-antimycotic solution (Thermo Fisher). Cultures were incubated at 37°C in a humidified atmosphere with 5% CO_2_.

### Infection of HeLa cells

On the day of infection, an overnight culture of *Salmonella* was diluted in LB media and allowed to grow until reaching late exponential phase. One day prior to infection, HeLa cells were transfected with pLAMP1-miRFP703 plasmid (Addgene #79998) using the NxT Electroporation system. Transfections were carried out using 10 μL tips containing 5 × 10^4^ HeLa cells and 1 μg of plasmid in R buffer (2 Pulses, 1005V, 35 ms). Transfected cells were then seeded onto Sarstedt (Germany) 8-well chambers (5 × 10^4^ cells per well) and cells were infected with *Salmonella* the following day at a multiplicity of infection (MOI) of 100. The infection by *Salmonella* was allowed to proceed for 1 h, after which cells were washed with DPBS and extracellular bacteria were eliminated by treatment with gentamicin (100 μg/mL) for 1 h. Infected HeLa cells were maintained in phenol-free DMEM containing 20 μg/mL of gentamicin.

### Bacterial biofilms for FLIM imaging

Bacterial cultures harboring pFPV::mcherryTYG were induced to form biofilms following a method adapted from Mon *et al*.^[25, 26]^ In brief, the bacterial cultures originated from a single colony in Luria broth (LB) and were left to incubate overnight at 37°C with agitation at 250 rpm. To initiate biofilm formation, 2 μL of the overnight culture were introduced into 198 μL of LB medium without salt in an 8-well glass bottom Ibidi μ-slide (Catalog No. 80806) and kept at 30°C for 2 days. Any LB growth medium without salt, which contained planktonic bacteria, was then removed from the preestablished biofilms, followed by washing, and replaced with pre-conditioned medium as needed.

### Infection of Zebrafish (*Danio rerio*)

Zebrafish husbandry and the static immersion of zebrafish larvae followed a protocol adapted from Desai *et al*.^[27]^ Initially, zebrafish adults and larvae were housed at our dedicated satellite facility. Eggs were obtained through a natural spawning protocol^[28]^ and incubated in embryo medium containing 0.001% methylene blue for 48 h at 28°C. Subsequently, the larvae were transferred to embryo medium supplemented with 25 μg/ml gentamicin for the following 6 hours before being moved to sterile embryo medium at 28°C. Fresh embryo medium was replenished daily. For static infections with *Salmonella* carrying pFPV::mcherryTYG (wild type and the *ssrB* null strain, Δ*ssrB*), 1 ml of an overnight culture grown in LB was inoculated into 10 ml LB broth supplemented with 100 μg/ml ampicillin and cultured for an additional 4.5 hours at 250 rpm at 37°C. The culture density was normalized by measuring the optical density at 600 nm. The bacterial culture was then harvested by centrifugation at 4,200 x g for 15 minutes at room temperature and resuspended in 1000 μl sterile embryo medium. Ten larvae at 5 days post-fertilization (dpf) were added to 8 ml of embryo medium in a 6-well polystyrene plate, followed by the addition of 80 μl of harvested bacterial cultures (10^9^ cfu/ml). Larvae exposed to an equal volume of PBS served as controls. After 24 h, the larvae were transferred to fresh embryo medium in a 6-well plate, with the medium being replaced daily. At 2 days post-infection, FLIM imaging was performed on infected fish.

### Infection of *C. elegans*

The *C. elegans* strain RZB286 was regularly cultured on NGM plates seeded with *E. coli* OP50 at 15°C. For infection, synchronized L4 nematodes were shifted to 25°C and exposed to *Salmonella* pFPV-mcherryTYG, spotted onto an NGM plate, for 24 h. Infected worms were then transferred to NGM plates to feed on *E. coli* for the duration of the experiment at 25°C.^[29]^ For imaging purposes, at day 6 following infection, the worms were immobilized using 25 mM Levamisole and embedded in 0.8% Ultrapure Agarose within an IBIDI chamber.

### FLIM imaging and data analysis

Fluorescence lifetime imaging was conducted using a TCS-SP8 3X system (Weltzar, Germany) equipped with a FLIM module. The mCherryTYG fluorophore was excited with a tunable pulsed white light laser (Leica WLL) operating at 20 MHz, with wavelengths ranging from 470 to 670 nm. Emission spectra were collected in the 580-720 nm range through the built-in acousto-beam splitter. Photon collection lasted approximately 0.3 to 1 minute using a HyD detector. Images typically had dimensions of 256 x 256 pixels, a scan speed of 400 Hz (lines per second), and a pinhole aperture set to 1.0 Airy units. The system was equipped with both HC PL APO CS2 100×/1.40 oil immersion and Plan Apochromat 20× oil objectives. All images, except those of zebrafish (which utilized the Plan Apochromat 20× oil objective), were acquired with the HC PL APO CS2 100×/1.40 oil immersion objective. The lifetime histograms of the mCherryTYG fluorophore were fitted to a sum of two-exponential decay functions (eq. 1) and deconvoluted with the instrument response function (IRF), which was generated using a reference dye, Coumarin 30 through FLIMfit.^[30]^

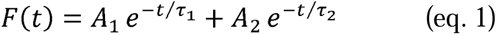

where, *A*_1_, *A*_2_ and *τ*_1_, *τ*_1_represents the amplitudes and time components of each decay, respectively. We then calculated the weighted average life-time (*τ_A_*) as:

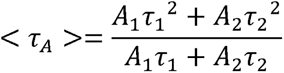

For image reconstruction, images were analysed using a 3×3 mean smoothing kernel applied via convolution to each image representing the various time bins, which helped minimize errors in local parameter estimates. TIFF files containing τ values were created using FLIMfit with 2 × 2 spatial binning before being convoluted with the fluorescence intensities. During FLIM imaging of bacterial infections, live cells were incubated at 37°C, 5% CO_2_ on the microscope, which was enclosed in a live cell imaging chamber (TOKAI HIT Microscope Stage Top Incubator).

## RESULTS AND DISCUSSION

### Characterization of mCherryTYG as a FLIM pH sensor

mCherryTYG is a pH-sensitive mutant of mCherry, where the methionine in the chromophore is replaced with threonine (M66T).^[23]^ With a pKa value of 6.8 and a broad dynamic range, mCherryTYG was selected as a potential FLIM-based pH sensor to unravel intricate pH-dependent processes in bacteria. To evaluate its functionality, we characterized its pH sensitivity both *in vitro* using purified protein solutions and *in vivo* within live bacterial cells through FLIM. *In vitro* characterization demonstrated that mCherryTYG exhibited a robust response to pH changes. Fluorescence microscopy of purified mCherryTYG revealed a tenfold decrease in brightness due to acid quenching as the pH shifted from 8.5 to 5.5 (Figure S-2A). This quenching was accompanied by a significant change in fluorescence lifetime, as confirmed by FLIM analysis (Figure 1). As expected, the fluorescence lifetime decay profile and FLIM image of purified mCherryTYG revealed a clear pH-dependent variation in fluorescence lifetime (Figure 1), as the pH was incrementally adjusted from 5.5 to 8.5 in 0.5-unit steps. The fluorescence lifetime varied from 1.06 ns at pH 5.5 to 2.15 ns at pH 8.5 (Figure 1), displaying an exceptionally broad dynamic range, consistent with previous findings.^[23]^ These results validated mCherryTYG as a reliable pH sensor with distinct, pH-dependent fluorescence lifetime variations.

**Figure 1.**
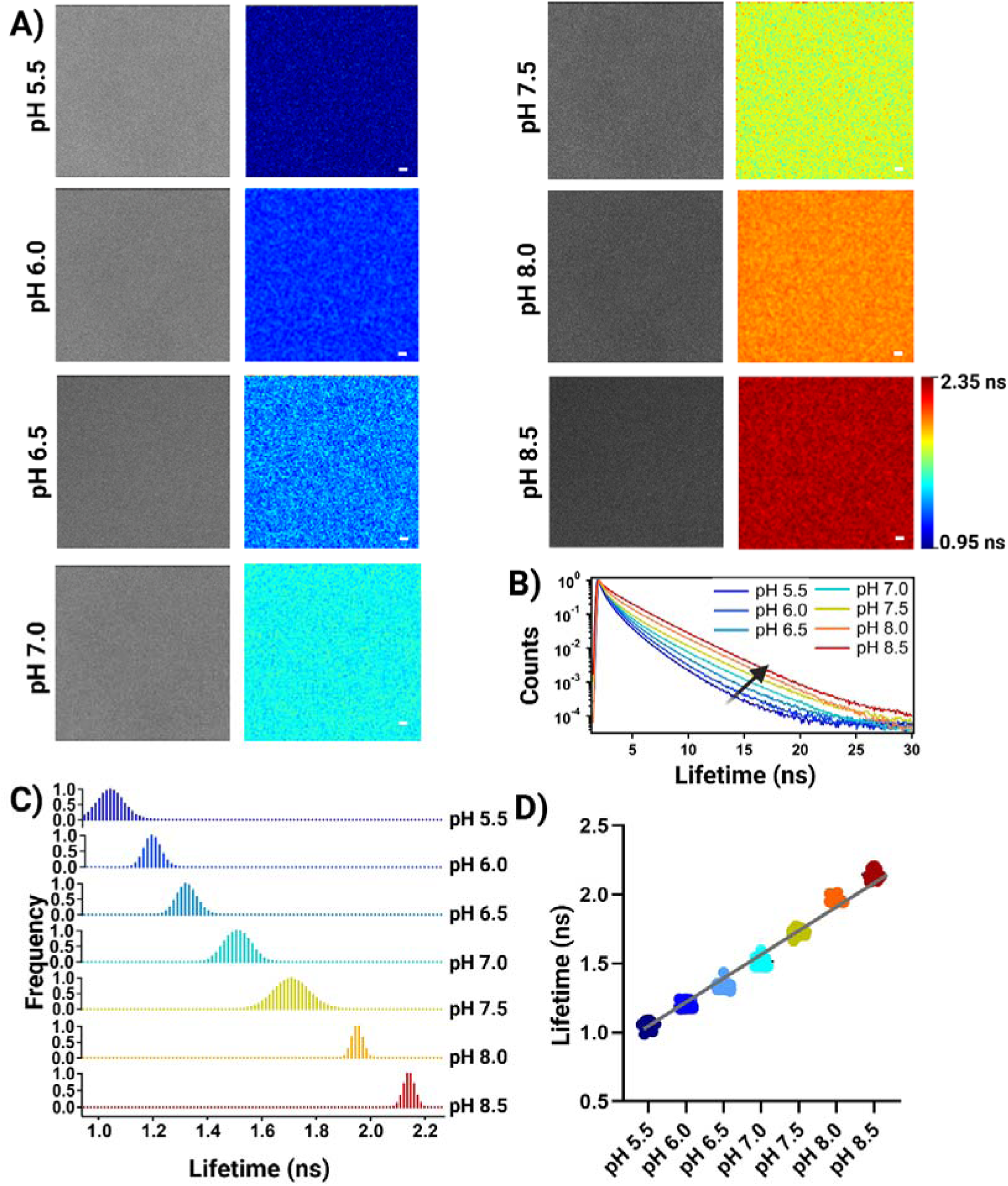
Fluorescence Lifetime Imaging (FLIM) of the purified mCherryTYG. (A) FLIM image of 10 μM mCherryTYG in calibration buffers of specific pH. The intensity image (left column) was convoluted with the fluorescence lifetime value for each pixel and then pseudo-colored (right column) using FLIMfit. Scale bars, 2 μm (B) Representative fluorescence lifetime histograms of mCherryTYG in calibration buffers of panel (A). The arrow points the lifetime as a function of pH. (C) Average lifetime histograms from the images of panel (A). (D) 30 regions of interest (each approximately 10 × 10 μm of the imaged area) were chosen for each pH buffer. These selections were made from three independent measurements, and the average lifetime τ was calculated. This panel also illustrates the pH-dependence of mCherryTYG in defined pH calibration buffers, from panel (A).

Building on these findings, we assessed the pH-dependent lifetime of mCherryTYG in live *Salmonella* Typhimurium cells. The sensor was expressed in the cytosol of live bacteria, and its lifetime response was measured under controlled conditions. To equilibrate external pH (pHe) with internal pH (pHi), we used the ionophore nigericin as a clamping agent, based on previous studies that demonstrated the ionophore nigericin was more effective at equilibrating external pHe with internal pHi than the weak clamping agent sodium benzoate.^[24]^ This approach revealed a nearly two-fold increase in fluorescence lifetime, from 0. 0.97 ns at pH 5.5 to 2.18 ns at pH 8.5 (Figure 2), almost matching the dynamic range observed *in vitro* (Figure 1). We observed an overall change of approximately 1.21 ns in the lifetime of mCherryTYG as the pH shifted from 5.5 to 8.5. The changes followed a linear trend, with a lifetime variation of about 0.40 ns per pH unit (Figure 2). Despite the quenching of fluorescence intensity at pH 5.5 (Figure S-2A), FLIM measurements remained readily observable. While various fluorescent proteins have shown pH-dependent lifetimes, ^[22, 23, 31–33]^ many like mScarlet, or E^2^GFP, exhibit different multi-exponential decays at varying pH levels, complicating decay fitting.^[22, 32]^ Ideally, FLIM data should fit the same decay model under all conditions. Unlike other pH-sensing proteins, the FLIM data from mCherryTYG can be consistently fit with the same decay model across all pH conditions tested. This uniformity simplifies data analysis and enhances the reliability of the sensor for FLIM-based applications.

**Figure 2.**
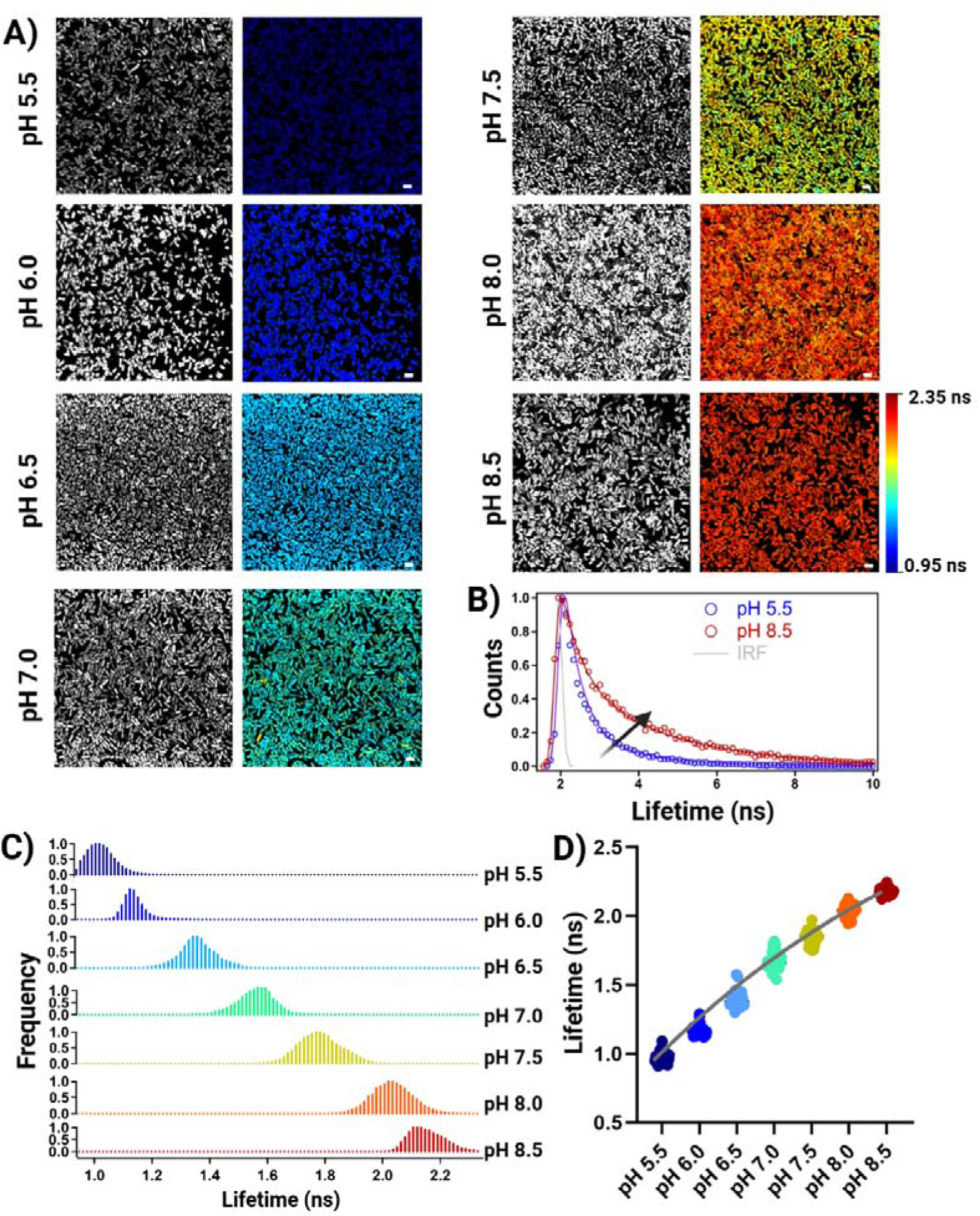
Calibration of mCherryTYG by FLIM in *Salmonella* Typhimurium cells expressing mcherryTYG. (A) The FLIM images depict *Salmonella* Typhimurium cells expressing mCherryTYG in specific calibration buffers. The intensity image (left column) was combined with the fluorescent lifetime value for each pixel and then pseudo-colored (right column) using FLIMfit. Scale bars, 2 μm. (B) Representative fluorescence lifetime histograms of mCherryTYG in solutions with pH 5.5 or pH 8.5. The fits were achieved with two-exponential decay functions combined with the instrumental response function (IRF). The graphs are normalized to the highest photon counts. (C) Lifetime histograms from the images of panel (A). (D) pH dependence of *Salmonella* Typhimurium cells expressing mCherryTYG in defined pH calibration buffers. The data was derived from the images from three independent experiments. The number of cells (N) ranged from 30 to 60.

Since mCherry is relatively inert, changes in lifetime due to non-specific interactions between mCherryTYG and the crowded cellular environment can be ruled out.^[34]^ This is supported by the observation that mCherryTYG maintained a pKa of 6.78, even in the crowded environment of live bacterial cells (see Figure S-3D). To further validate the robustness of mCherryTYG, we assessed its performance under various experimental conditions. While fluorescence lifetime is more resilient than fluorescence intensity, it can still be influenced by factors other than pH (see Figure S-2 and S-3). We observed no significant changes in the lifetime response in different buffers, including phosphate-buffered sodium, MES, and MOPS buffers containing Tris and Bis-Tris (Figure S-2C). Additionally, the lifetime of mCherryTYG remained unchanged with various concentrations and laser power (Figure S-2B and S-3A). A previous study also showed that mCherryTYG exhibited no sensitivity to differences in physiological salt or differences in the redox environment.^[23]^ Temperature can affect fluorescence lifetimes, but we found no changes in mCherryTYG lifetimes within the physiological pH range across the tested temperatures 23°C - 37°C (Figure S-3C). However, a minor decrease in lifetime was observed at higher pH levels (8.0–8.5), but this did not compromise the overall performance of the sensor (Figure S-2 and S-3). Collectively, these results establish mCherryTYG as a highly sensitive and reliable FLIM-based pH sensor. Its ability to accurately track intracellular pH changes under various conditions makes it an invaluable tool for exploring the pH-dependent mechanisms underlying bacterial pathogenesis, adaptation, and lifestyle transitions, particularly in *Salmonella* Typhimurium.

### Mapping pH dynamics inside *Salmonella in-vitro* and in heterologous hosts

#### Imaging pH heterogeneity in bacteria in vitro

Gram-negative bacteria such as *E. coli* and *Salmonella* Typhimurium have long been considered neutralophilic, maintaining their intracellular pH between 7.2 and 7.8.^[35]^ More recent studies have challenged this notion, reporting that bacteria acidify in response to changes in pHe.^[9,21]^ We used our mCherry TYG FLIM sensor to measure the cytoplasmic pH of *Salmonella* Typhimurium in various environmental conditions: LB at pH 4.5, MgM at pH 5.6, and MgM at pH 7.2 (Figure 3). Our findings revealed that the cytoplasmic pH was ∼ 6.80 at an extracellular pH (pHe) of 7.2, and 6.04 at pHe 5.6, indicating a relatively homogeneous lifetime (Figure 3). These observations align with previous pH measurements obtained using the ratiometric indicator BCECF-AM^[21]^ and a FRET DNA biosensor, the I-switch^[9]^. This result indicates that previous heterogeneity observed with the pH-sensitive GFP (pHluorin) likely stems from both variations in plasmid expression and the clamping method.^[21]^ The measured cytoplasmic pH of 5.78 at pHe 4.5 was comparable to the pH of 6.04 observed at pHe 5.6, indicating that either mCherryTYG exhibits reduced sensitivity at lower pH levels, or there is a limit to cytoplasmic acidification. Overall, our results reinforce the notion of a sustained decrease in bacterial cytoplasmic pH in response to acid stress.

**Figure 3.**
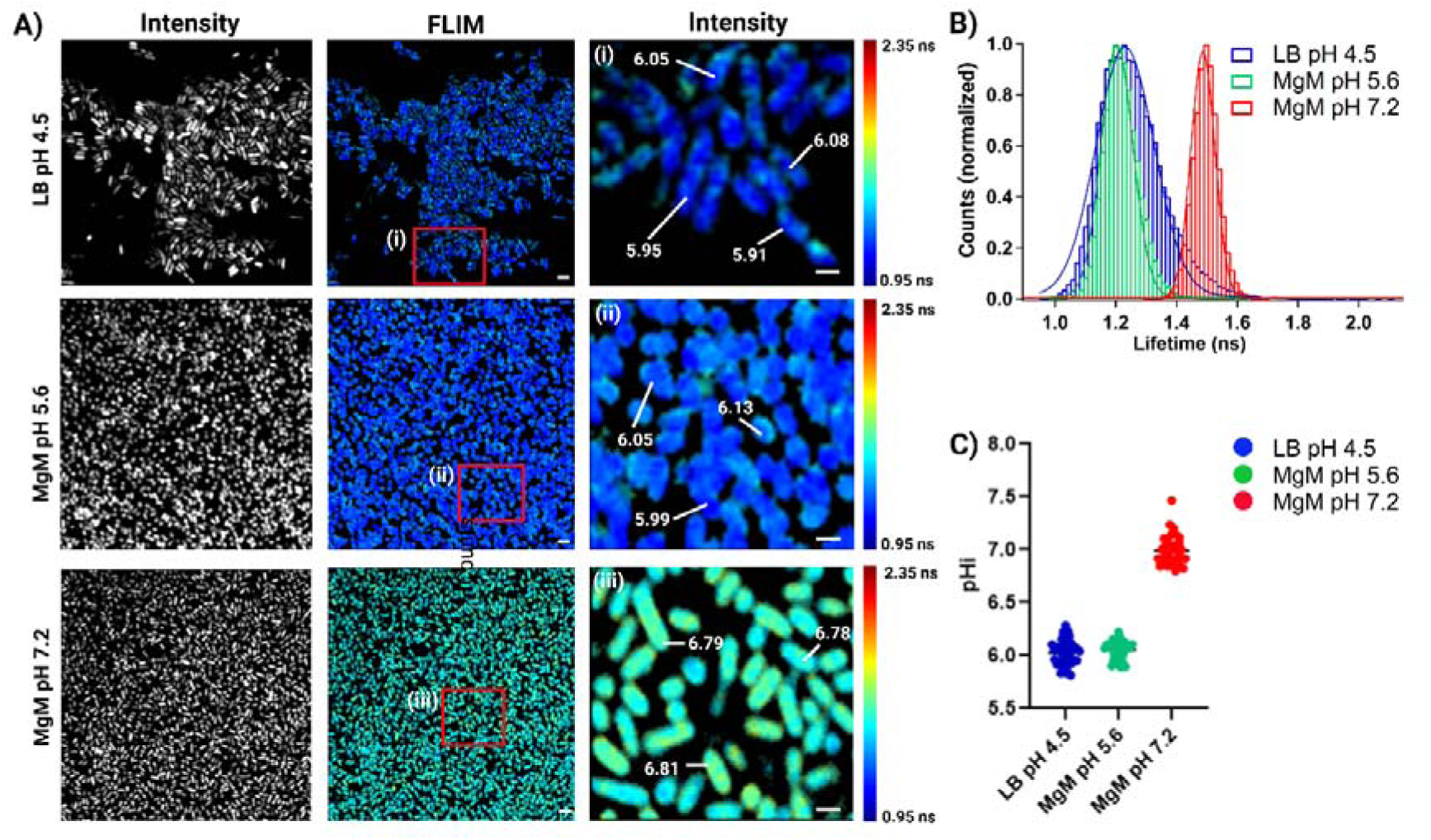
FLIM images of *Salmonella* under varying environmental conditions. (A) The FLIM images show *Salmonella* Typhimurium cells expressing mCherryTYG in LB pH 4.5, MgM pH 5.6, and MgM pH 7.2. The intensity image (left column) combines the fluorescent lifetime value for each pixel and is pseudo-colored using FLIMfit. Insets on the right display expanded views with pH values of selected bacteria indicated. (B) Representative fluorescence lifetime histograms of mCherryTYG in *Salmonella* in LB pH 4.5, MgM pH 5.6, and MgM pH 7.2. (C) Quantification of pH values from panel A. Scale bars, 2 μm, inset; 1 μm.

The high sensitivity of FLIM-based measurements of mCherryTYG pH revealed a slight localized heterogeneity in pH within bacterial cells. Specifically, the pH near the membrane region was slightly higher than in the core area of the bacteria, a distinction that other methods have failed to visualize (Figure 3 insets).

#### Bacterial pH dynamics during infection in HeLa cells

Once *Salmonella* enters a cell, it resides within the *Salmonella*-containing vacuole (SCV), which is acidic, while some bacteria escape to the neutral cytoplasm. These two subpopulations experience vastly different intracellular pH levels, which significantly influences their physiological states and virulence programs. Acidified bacteria within the SCV exploit the acidic pH to activate a type III secretion system (T3SS), a critical secretory apparatus responsible for manipulating host cell processes. It was therefore of interest to measure the pH sensed by *Salmonella* both inside and outside of the SCV, and to investigate any heterogeneity of bacterial pH inside mammalian cells.

To measure the pH of *Salmonella* both inside and outside the SCV, we transfected HeLa cells with *p*lamp1-miRFP670 as a marker for the SCV and endosomal tubulation. Using both Lamp1-miRFP670 and mCherryTYG allowed us to distinguish whether *Salmonella* was located inside or outside the SCV (Figure 4). Four hours post-infection, we collected confocal and FLIM images of mCherryTYG-expressing *Salmonella* in HeLa cells (Figure 4A). *Salmonella* residing within the SCV (defined as co-localized with Lamp1) are acidic (∼pH 5.89), while bacteria outside the SCV remain neutral (pH 7.10). This finding was supported by the dual lifetime distribution shown in Figure 4C-D. Outside the SCV, we observed little to no heterogeneity in fluorescence lifetime. After 16 hours post-infection, most bacteria remained within an SCV-like structure (see Figure S-4A-B). Within what appears to be the same SCV, only a small fraction of bacteria were acidified (∼pH 5.94), while the majority remained neutral (pH 7.33), as illustrated in Figure 4B-C. The dual lifetime distribution results in Figure 4C indicated that less than ∼17% of the bacterial population were acidified (exhibiting a lower lifetime), highlighting a phenotypic heterogeneity in terms of the pH of the bacteria inside host cells. This was in marked contrast to the homogeneous acidification observed in acid inducing conditions *in-vitro* (Figure 3). These findings highlight the heretofore unreported dynamic and heterogeneous nature of the intracellular niches of *Salmonella* and its reliance on precise pH regulation to optimize its virulence strategy during infection, an observation made possible by the utility of the mCherryTYG FLIM sensor.

**Figure 4.**
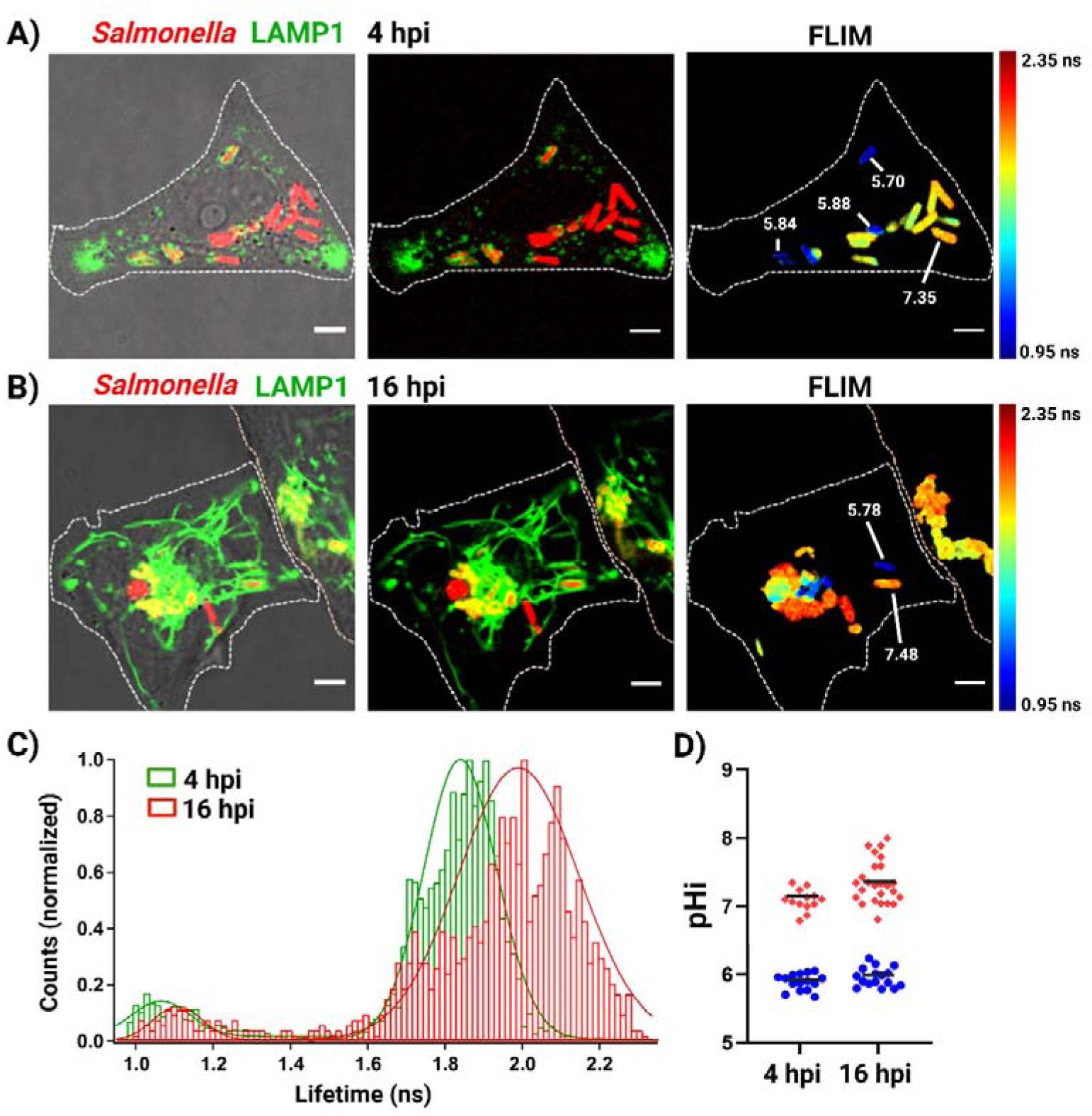
Tracking the pH of *Salmonella* during HeLa cell infection. (A) HeLa cells harboring Lamp1-miRFP703 were infected with *Salmonella* (red) and imaged using confocal microscopy (left and middle panels) 4 hours post-infection. The right panel shows the corresponding FLIM image. (B) The left and middle panels show a confocal image of HeLa cells infected with *Salmonella* Typhimurium expressing mCherryTYG, 16h post-infection. The right panel displays the corresponding FLIM image, with the pHi of selected bacteria indicated. (C) Representative fluorescence lifetime histograms of mCherryTYG in *Salmonella* during HeLa cell infection. (D) Quantification of pH values from panels A and B (insets). Scale bar, 5 μm.

### Three-dimensional mapping of bacterial pH in *Salmonella* biofilms

Under acidic pH, *Salmonella* Typhimurium activates virulence genes for invasion and systemic infection, while neutral pH triggers dormancy and biofilm formation. Forming a biofilm enables immune evasion and persistence as an asymptomatic carrier. Biofilms act as protective niches, allowing the bacteria to remain dormant and avoid immune clearance, while still being able to reactivate under favorable conditions when the host immune system is compromised or when environmental conditions shift. This switch between active virulence and dormancy is closely tied to pH changes within the host, highlighting the critical role that pH plays in regulating bacterial behavior and persistence. ^[12, 27, 29]^

To better understand the pH dynamics within *Salmonella* biofilms, cultures were grown in LB medium without salt. Fluorescence and FLIM images were captured at various Z-stack depths to visualize the three-dimensional structure and pH variations within the biofilm (Figure 5A-C). The 3D fluorescence projection in Figure 5A, color-coded by depth, highlights the architecture of the *Salmonella* biofilm. The corresponding FLIM images in Figure 5C were taken at three distinct depths: bottom, middle, and top of the biofilm, showing the pH detected by a few of individual bacteria at each stack. These images illustrate a clear lifetime (pH) gradient across the biofilm, with a more acidic environment at the bottom and a neutral or slightly basic pH at the surface. The lifetime distribution histograms in Figure 5D further emphasize the variation in fluorescence lifetime along the Z-axis, reflecting differences throughout the biofilm thickness. The average pH per Z-stack at the bottom of the biofilm was approximately 6.45, and increased to 7.4 at the top of the biofilm, spanning a thickness of 8 µm from the bottom to the surface (Figure 5C-E). These pH measurements support the existence of a stratified pH profile within *Salmonella* biofilms, with a slightly acidic bottom and a more alkaline surface, similar to the pH gradients observed in other bacterial biofilms, including *E. coli*^[36]^ and *Pseudomonas aueruginosa*^[37]^. Interestingly, the pH of individual *Salmonella* cells within the same stack also varied, suggesting that local microenvironments within the biofilm contributed to the observed pH heterogeneity. A 3D projection movie (Supplementary Movie S1) of *Salmonella* biofilms reveals a dynamic lifetime (pH) gradient, with fluorescence lifetimes increasing from the bottom to the top of the biofilm, corresponding to a rise in pH from acidic to neutral (Figure 5E). These findings further confirm that *Salmonella* biofilms, like those formed by *E. coli* and *Pseudomonas aeruginosa*, display a pH gradient across their thickness, with a more acidic core, bottom and a neutral or basic surface. This pH stratification within biofilms provides a dynamic environment that not only facilitates bacterial survival but also helps the bacteria adapt to changing conditions. The acidic core is likely involved in protecting the bacteria from host immune responses and antimicrobial treatments, while the alkaline surface aids in nutrient uptake and persistence in the host.^[36, 38, 39]^ The pH gradient within biofilms is a critical factor in the ability of *Salmonella* to survive in diverse host environments, resist immune responses, and contribute to the chronicity and recalcitrance of infections.^[36, 38, 39]^ These findings highlight the importance of targeting the pH microenvironments of biofilms as a potential strategy for combating persistent bacterial infections and improving treatment outcomes.

**Figure 5.**
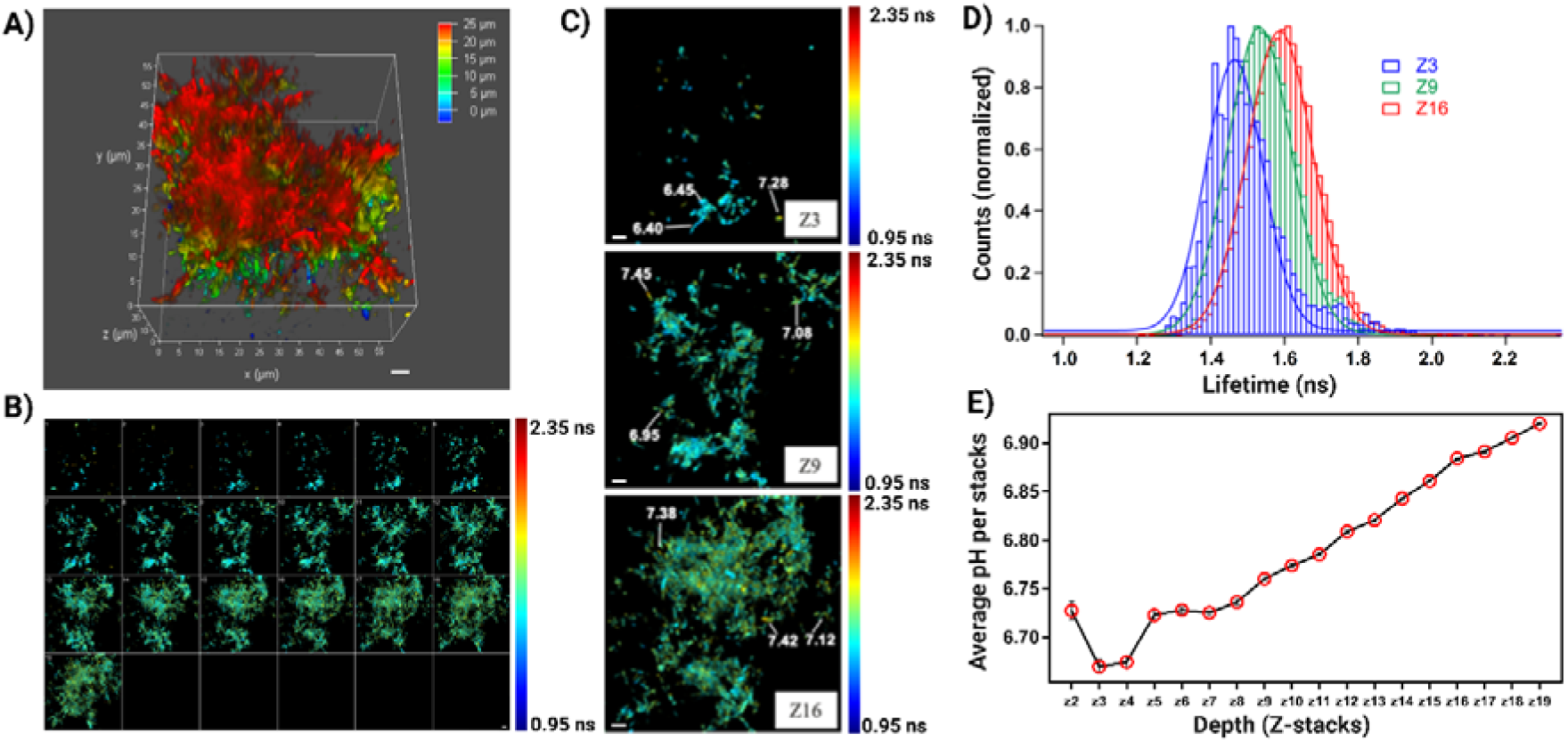
3D pH mapping of *Salmonella* biofilms. (A) 3D projection of fluorescence microscopy image of a *Salmonella* biofilm. Scale bars, 10 μm. (B) Corresponding FLIM image at Z-stacks spanning 0.5 µm across the biofilm. Scale bars, 10 μm. (C) FLIM image at three selected Z-positions (bottom, middle, and surface), with the pH of selected bacteria indicated by arrows. Scale bars, 10 μm. (D) Corresponding lifetime histogram from panel C. (E) Quantification of average pH values per Z-stack from the bottom to the top of the biofilm.

#### pH mapping of Salmonella inside heterologous hosts

The intestinal lumen of eukaryotic systems serves as a microenvironment where bacterial colonization occurs and factors such as pH can affect bacterial growth and biofilm formation, both of which are critical to bacterial pathogenesis. *C. elegans* is a simple, transparent eukaryotic model that is useful for studying bacterial infection lifestyles. When *C. elegans* was fed *Salmonella* in place of *E. coli,* its native food source, biofilms formed in the intestinal lumen. Biofilm formation was advantageous to the worm, expanding longevity through the activation of a mitogen-activated protein kinase pathway.^[12,39]^ To quantify the pH experienced by *Salmonella* throughout the worm, L4 stage (adult) nematodes were exposed to wild type *Salmonella* Typhimurium for 24 h, allowing sufficient time for bacterial colonization and adaptation. Figure 6 shows representative FLIM images of a nematode infected with *Salmonella* expressing mCherryTYG, segmented into three regions. Clear lifetime differences across the three segmented regions of the worm intestinal lumen reflected distinct pH conditions experienced by *Salmonella*. The pH ranged from ∼7.10 in the head region to ∼6.45 towards the tail (Figure 6B-C), indicating a gradient that influences bacterial physiology and biofilm formation. Indeed, biofilms were identified in the anterior regions of *C. elegans*, where the pH is neutral (Figure 6).^[37]^ This pH gradient identified here was also supported by previous observations using pH nanosensors,^[40]^ further validating the reliability of our FLIM-based mCherryTYG sensor. These findings demonstrate the power of combining *C. elegans* as a model system with advanced imaging technologies to unravel the complex interplay between host microenvironments and bacterial pathogenesis.

**Figure 6.**
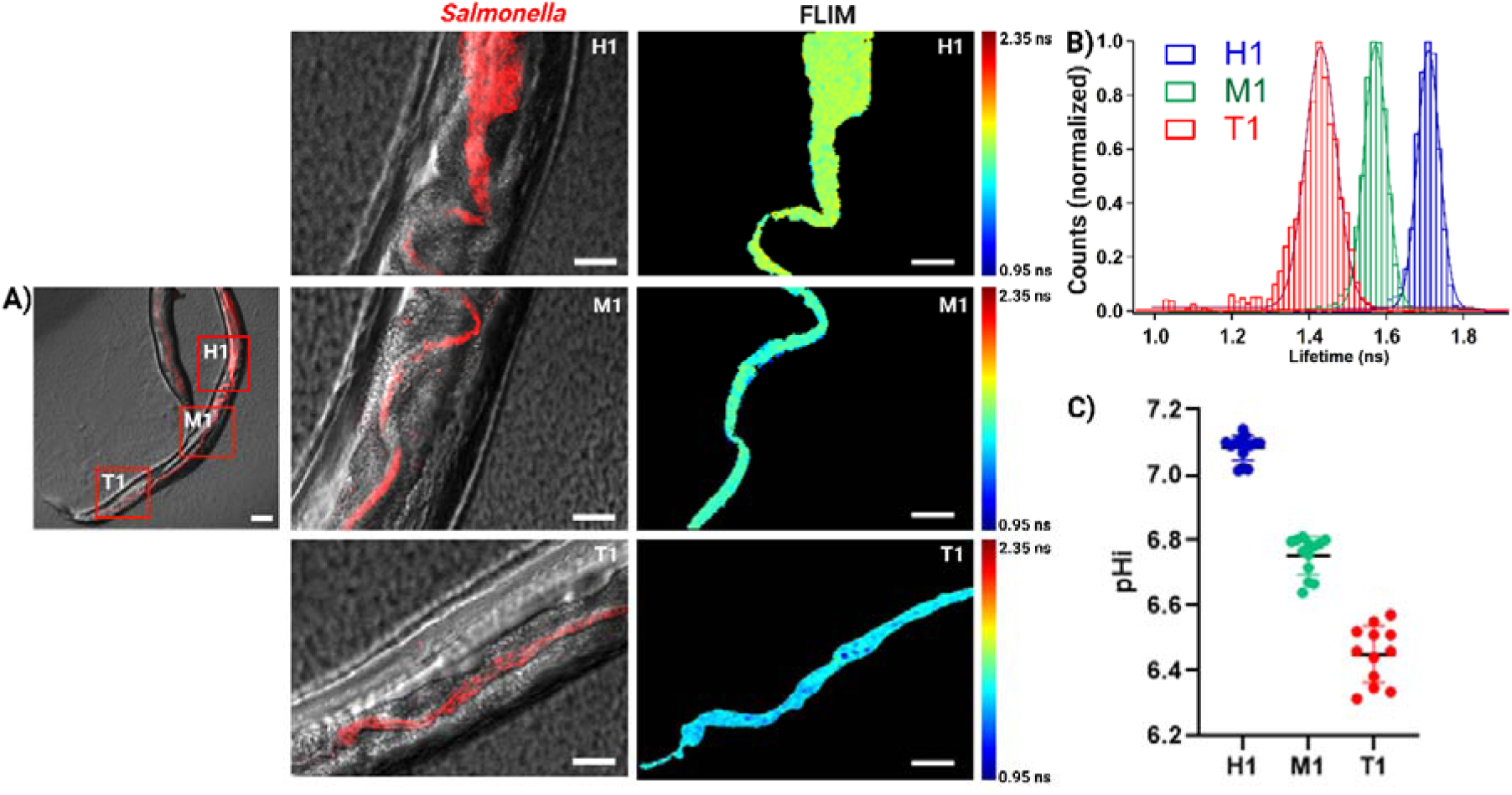
Mapping the pH of *Salmonella* in *C. elegans*. (A) Left and middle panels display merged fluorescence microscopy images of *C. elegans* infected with *Salmonella* expressing mCherryTYG, highlighting three distinct regions along the worm. The right panel shows the corresponding FLIM image. Scale bars, 100 μm, inset; 20 μm. (B) Representative fluorescence lifetime histograms of *C. elegans* from the insets in panel A. (C) Quantification of pH values across the three regions, labelled H1, M1, and T1.

The zebrafish model closely mirrors human systems in developing immunity against infectious microbes ^[41]^ Among its many attributes, the zebrafish gut contains a specialized region of acidic Lysosome-Rich Enterocytes (LRE) that play a crucial role in intestinal homeostasis.^[42–44]^ This region is essential for protein endocytosis from the intestinal lumen and bacterial uptake, making it a key site for investigating microbial infections and their interaction with the host immune system.^[32, 33, 45]^ Traditionally, the luminal pH in zebrafish larvae has been measured using pH-sensitive dyes such as m-cresol purple (0.2% in embryo media) and phenol red (2% in embryo media), which rely on comparison against calibrated pH standards ranging from pH 6 to 10. While these dyes are useful for providing rough estimates of pH, their resolution is limited, and they fail to capture subtle pH changes or localized microenvironmental variations within the intestine. Additionally, functional lysosomes in the LRE region have been assessed for acidic conditions using neutral red (2.5 µg/ml for live larvae) and alcian blue (0.01% for PFA-fixed larvae) dyes during *E. coli* infection studies.^[43]^ While neutral red provides a visual confirmation of acidic conditions within live larvae, it lacks the precision necessary for accurate pH quantification. As a result, the precise pH profile sensed by bacteria in the LRE region, as well as in the anterior regions of the zebrafish gut, remains poorly characterized. To address these limitations, we set out to accurately measure the pH detected by *Salmonella* Typhimurium in both the LRE and anterior regions of the zebrafish gut using FLIM and our pH-sensitive fluorophore mCherryTYG. To establish the baseline, the fluorescence lifetime in zebrafish fed with PBS was first assessed (see Figure 7B and Figure S-5B). The average background fluorescence lifetime was determined to be 4.44 ns, which was outside the dynamic range of the mCherryTYG lifetime for pH sensing (Figure 7E). Thus, the background fluorescence was unlikely to interfere with the subsequent pH measurements in zebrafish. Next, purified mCherryTYG protein was orally administered to zebrafish larvae, and FLIM images were captured two hours after ingestion. FLIM images revealed a significantly distinct mCherryTYG fluorescence lifetime in the LRE region, as highlighted in the segmented image inset in Figure 7C. The observed lifetime in the LRE was measured at 1.16 ns, corresponding to a pH of approximately 5.92 (Figure 7F-G). This finding suggests that the mCherryTYG protein that was taken up by the LREs was in an acidic lysosome. To investigate the pH sensed by *Salmonella* Typhimurium in the LREs, zebrafish larvae were infected with *Salmonella* expressing pFPV::mCherryTYG, and FLIM imaging was performed two days post-infection. FLIM images showed that the fluorescence lifetime of *Salmonella* localized in LREs was 1.12 ns, corresponding to a pH of approximately 5.84 (Figure 6E-G). This similarity in lifetimes and pH values between the mCherryTYG protein and *Salmonella*-containing mCherryTYG suggests that *Salmonella* was taken up by LREs, and residds in an acidic intracellular compartment. In contrast to the acidic environment in the LRE, *Salmonella* present in the anterior region of the zebrafish intestinal lumen experienced a more alkaline pH environment, with an average pH of 7.33 and a corresponding average lifetime of 1.78 ns (Figure 7E-G and see Figure S-5A). This marked difference in pH levels between the LRE and anterior regions highlights the spatial heterogeneity of the zebrafish intestinal microenvironment. Such variations likely play a critical role in shaping bacterial physiology, behavior, and adaptation during colonization and infection. Together, these findings provide a comprehensive understanding of the pH dynamics experienced by *Salmonella* in the zebrafish model, emphasizing the utility of FLIM and mCherryTYG for elucidating microenvironmental conditions critical to bacterial pathogenesis.

**Figure 7.**
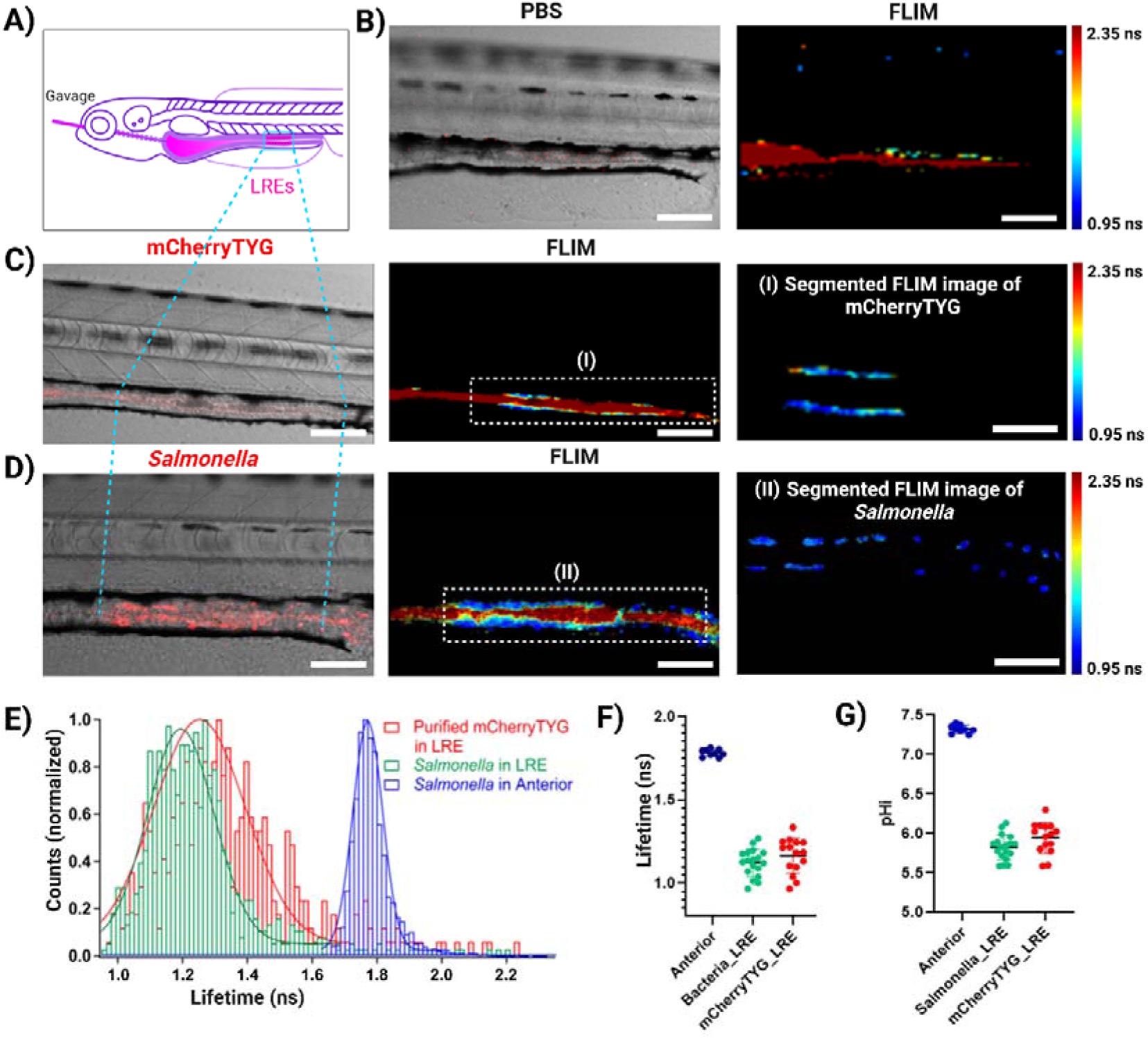
Mapping the pH of *Salmonella* in zebrafish. (A) Cartoon of zebrafish highlighting the LRE region. (B) Merged fluorescence microscopy images of zebrafish gavaged with PBS control. Right panel: FLIM image of zebrafish background. (C) Left panel: Merged fluorescence microscopy images of C. Zebrafish gavaged with purified mCherryTYG, with the corresponding FLIM image in the middle panel. Inset (I): Manually segmented FLIM image of mCherryTYG protein in LREs. Scale bars, 50 μm. (D) Merged fluorescence microscopy images of zebrafish infected with *Salmonella*, with the corresponding FLIM image in the middle panel. Inset (II): Manually segmented FLIM image of *Salmonella* in LREs. Scale bars, 50 μm. (E) Representative fluorescence lifetime histograms of purified mCherryTYG in LREs, *Salmonella* at LRE and anterior region from panel C, D and Figure S-5. (F-G) Quantification of lifetime and pH values of *Salmonella* at anterior and LRE regions compared to purified mCherryTYG at the LRE region.

## CONCLUSION

This study elucidates the critical role of intracellular pH dynamics in *Salmonella* Typhimurium pathogenesis and lifestyle adaptations, leveraging the innovative pH-sensitive fluorophore mCherryTYG in conjunction with FLIM. By integrating precise temporal and spatial imaging techniques, we provide a comprehensive understanding of bacterial pH regulation across diverse environments and infection contexts. We developed and validated mCherryTYG, a genetically engineered pH sensor, which demonstrated exceptional sensitivity across a broad pH range (5.5–8.5) with consistent fluorescence lifetime responses. Unlike traditional ratiometric sensors, mCherryTYG retained its reliability across varying physiological conditions, including different buffers, ionic environments, and temperature ranges. FLIM-based measurements established mCherryTYG as a robust and accurate platform for non-invasive quantification of intracellular pH, addressing limitations of conventional methods. Using this tool, we uncovered critical insights into pH regulation in *Salmonella*. *In vitro*, we observed sustained intracellular acidification in response to acidic environments, correlating with the activation of virulence. FLIM imaging revealed a uniform intracellular pH within *Salmonella* under inducing conditions *in vitro*, highlighting the precision and consistency of mCherryTYG over previous methods, which had suggested heterogeneity. In host infection models, we demonstrated distinct pH environments encountered by *Salmonella* during infection. In HeLa cells, we observed a dichotomy between acidified *Salmonella* residing within the SCV (∼pH 5.89) and neutral cytoplasmic bacteria (∼pH 7.10). Interestingly, at later stages of infection, phenotypic variability emerged within SCVs, with less than ∼17% of bacteria maintaining an acidic pH. These findings emphasized the role of pH as a critical signalling mechanism for virulence activation. In biofilms, FLIM imaging revealed a stratified pH profile, with an acidic bottom or core (∼pH 6.45) and neutral surfaces (∼pH 7.4), mirroring patterns observed in biofilms of other pathogens. This stratification likely contributes to bacterial persistence, immune evasion, and chronic infection resilience. In heterologous host models, such as *C. elegans* and zebrafish, FLIM imaging revealed dynamic pH gradients influencing bacterial adaptation to host specific niches. In *C. elegans*, *Salmonella* experienced a neutral pH (∼7.10) in the anterior where biofilms are formed, and progressively acidic conditions (∼6.45) toward the tail, reflecting the intestinal environment of the host. Similarly, in zebrafish, *Salmonella* encountered an acidic pH (∼5.84) in lysosome-rich enterocytes (LREs), but a more neutral pH (∼7.33) in the anterior intestine. These findings underscore how local pH gradients shape bacterial colonization strategies and persistence in host systems.

Overall, these findings establish mCherryTYG-FLIM as a transformative platform for studying bacterial pH regulation and its functional implications. The ability to track intracellular pH with high spatial and temporal resolution has revealed that pH serves as a master regulator of bacterial behavior, influencing virulence, adaptation, and persistence. The broader implications of this work extend beyond *Salmonella*. The versatile mCherryTYG-FLIM platform provides a foundation for investigating pH-dependent mechanisms in other pathogens and complex biological systems, paving the way for novel therapeutic interventions. By targeting pH-mediated pathways, our research offers innovative strategies to mitigate bacterial virulence, biofilm resilience, and chronic infections.

## ABBREVIATIONS

(FLIM): Fluorescence lifetime imaging microscopy
(MOI): multiplicity of infection
(TRFS): Time-resolved fluorescence spectroscopy
(SCV): *Salmonella*-containing vacuole
(LRE): Lysosome-Rich Enterocytes

## Acknowledgments

We are grateful to Dasvit Shetty for protein purification and for the exceptional expertise of the Mass Spectrometry Core at the University of Texas Medical Branch (UTMB).

## Author Contributions

MKS and LJK conceptualized and designed the research; MKS, MF and RD performed the experiments; PZ purified the protein; MKS analyzed the data, MKS and LJK wrote the original draft, reviewed and edited the manuscript. RD edited the references. All authors have read and agreed to the published version of the manuscript.

## Additional information

Supplementary information is available for this paper online.

## SUPPLEMENTARY FIGURES

**Figure S-1.**
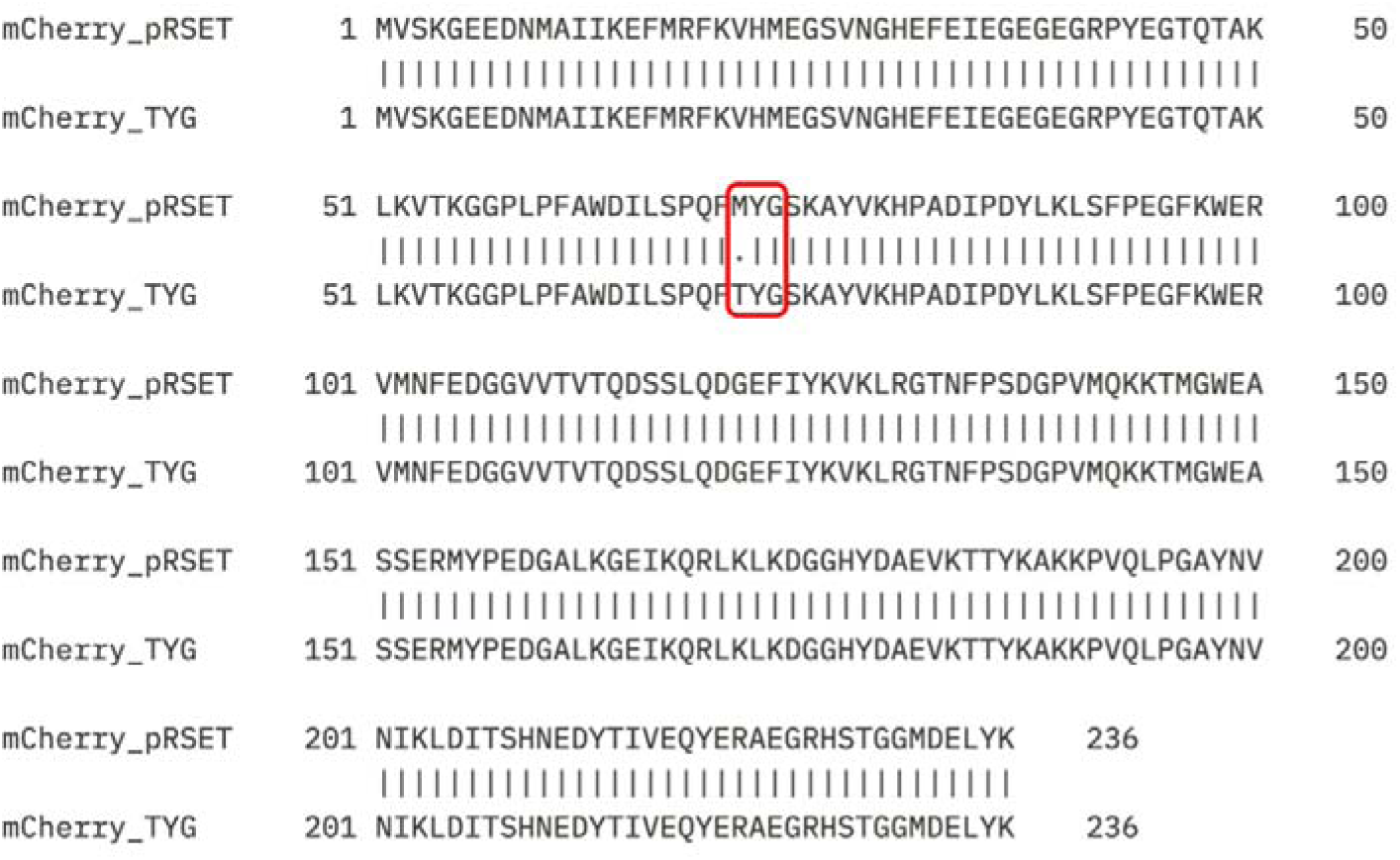
Multiple sequence alignment of mCherry protein using Emboss needle software. In pRSET-B-mCherry plasmid (addgene#108857), Methionine in position 66 of wild-type mCherry has been replaced with Threonine by site-directed mutagenesis, generating mCherryTYG (PDB6MZ3), a pH-sensitive mutant fluorophore.

**Figure S-2.**
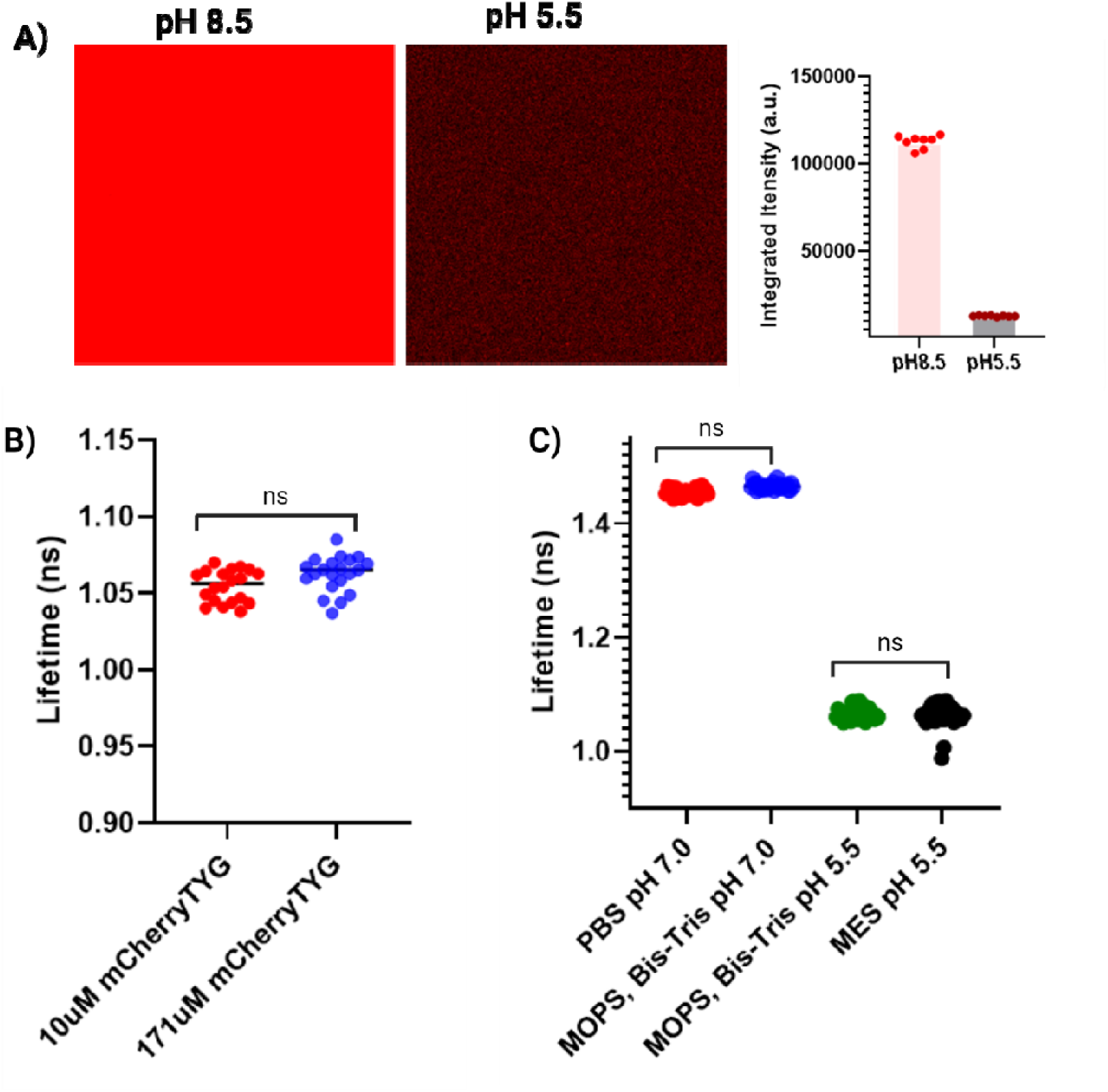
Examining the characteristics of mCherryTYG. (A) A representative confocal image of purified mCherryTYG is displayed, taken at the same laser power but at varying pH levels. The right panel quantifies the fluorescent intensity at two different pH levels, showing an approximate ten-fold increase in intensity when the pH shifts from 5.0 to 8.5. (B) No change in fluorescence lifetime was observed in MES at pH 5.5 as mCherryTYG concentration varied from 10 µm to 171 µm. The lifetime of mCherryTYG was not influenced by its concentration. This was confirmed by a NOVA test with Tukey’s (HSD) post hoc, which shows no significant difference. The data was derived from the images from three independent experiments (C) The lifetime of mCherryTYG remains constant regardless of the buffer composition, but varies with different pH levels. A NOVA test with Tukey’s (HSD) post hoc shows no significant difference. The data was derived from images from three independent experiments.

**Figure S-3.**
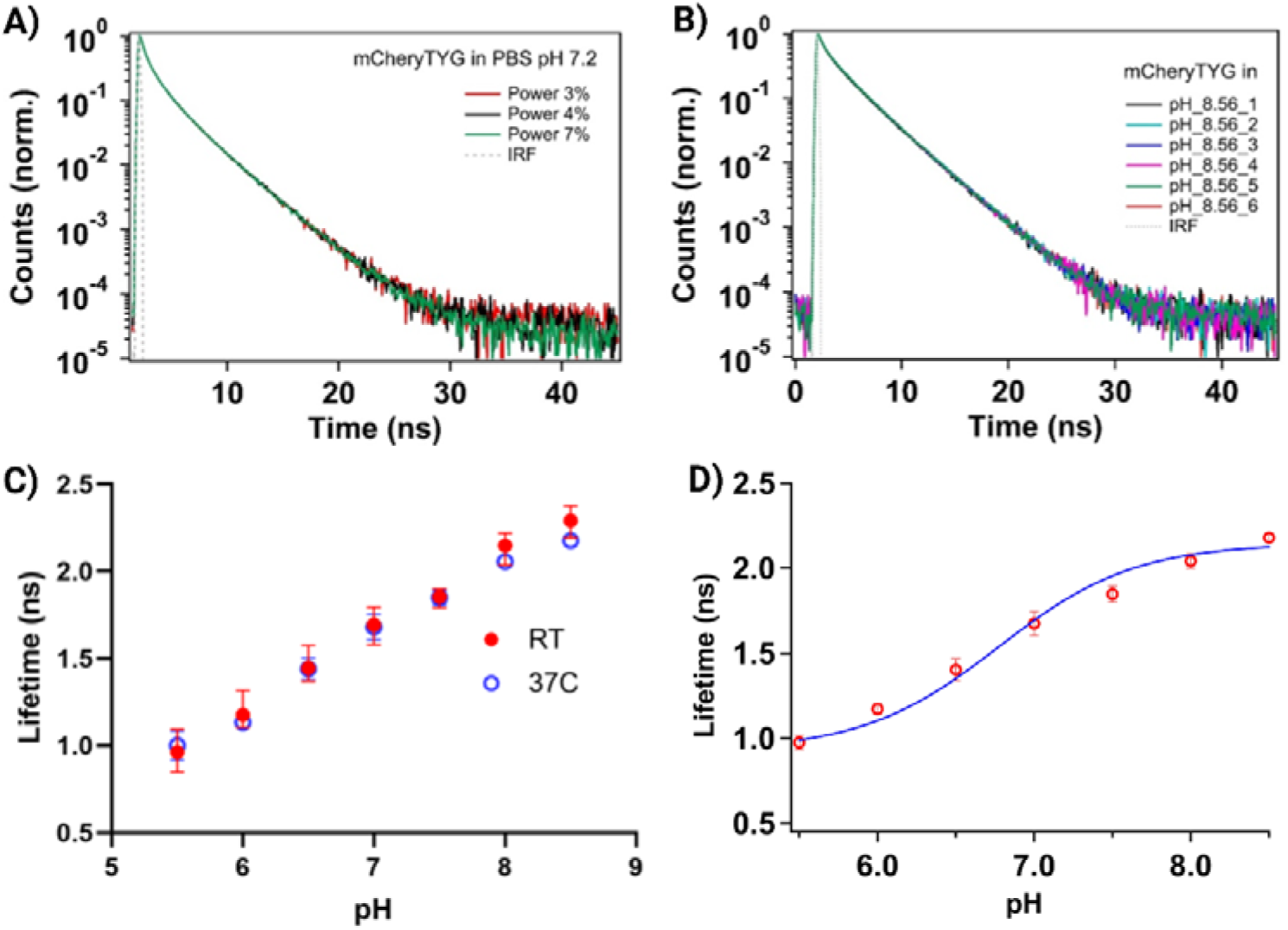
(A) mCherryTYG lifetime is independent of laser excitation. (B) Reproducibility of lifetime data collected at different times. (C) The response of mCherryTYG to pH changes at various temperatures. The graph illustrates the effect of temperature on the pH response of 10 µM mCherryTYG in a calibration buffer with a specified pH, showing that the lifetime response to pH is not significantly influenced by temperature. Each data point represents the average of three measurements taken on different days. (D) The graph of average lifetime (<τ>) versus pH for mCherryTYG expressed in *Salmonella* cells shows a calibration fit to the Henderson-Hasselbalch equation, revealing a pKa of ∼6.78. The data is based on images from three independent experiments.

**Figure S-4.**
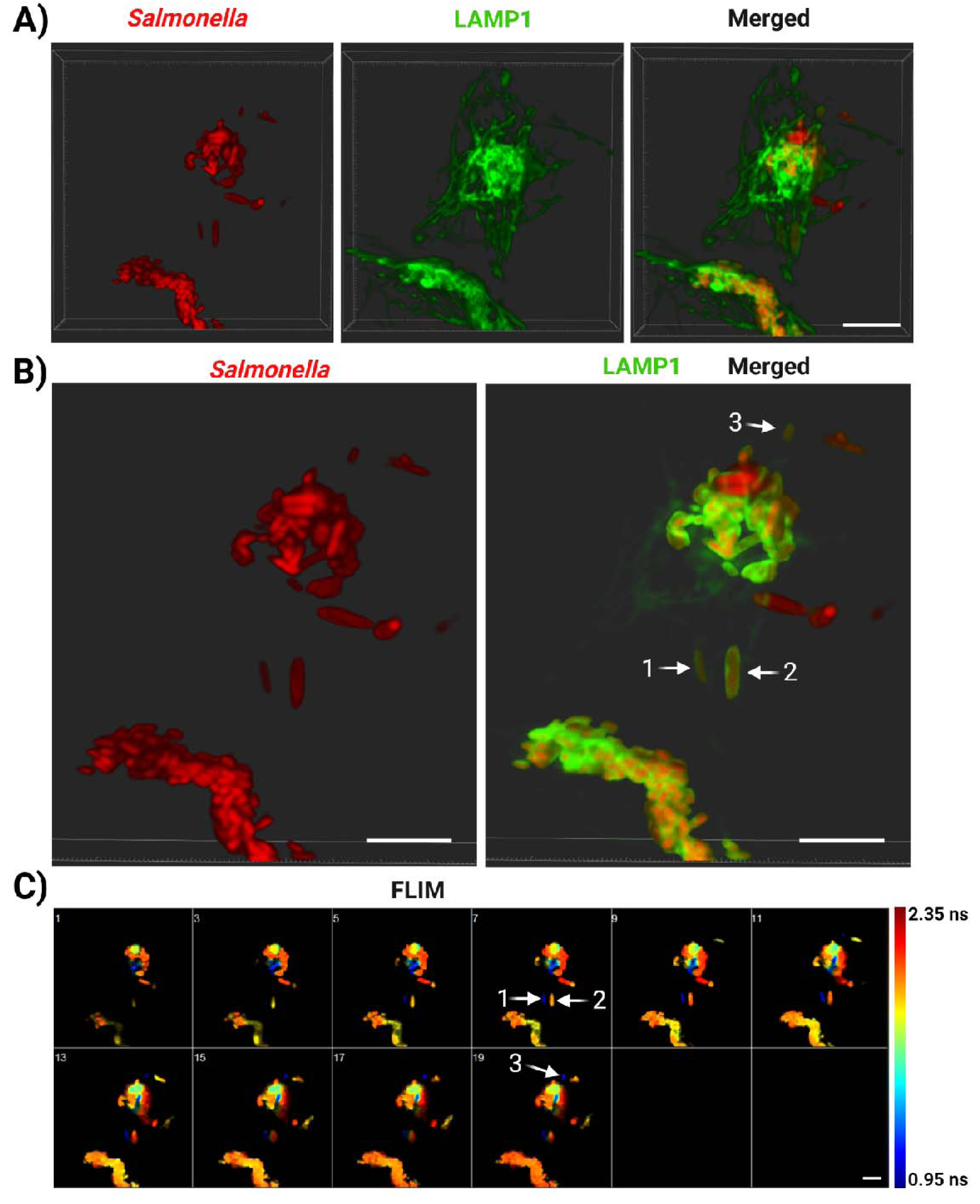
(A) Surface rendering of *Salmonella* and Lamp1 confocal images. (B) The same image is shown with adjusted opacity of the green channel (representing Lamp1) to enhance visualization of bacteria enclosed by Lamp1 within *Salmonella*-containing vacuoles (SCVs). The images clearly depict bacteria both within SCV-like structures and outside them. (C) Comparison of FLIM images with the corresponding regions in the right panel of (B). This comparison highlights that many bacteria within SCV-like structures are not acidified. For instance, three bacteria (labelled 1, 2, and 3) co-localize with Lamp1. FLIM data reveal that bacteria 1 and 3 were acidified, whereas 2 was not. Scale bars: 10 μm.

**Figure S-5.**
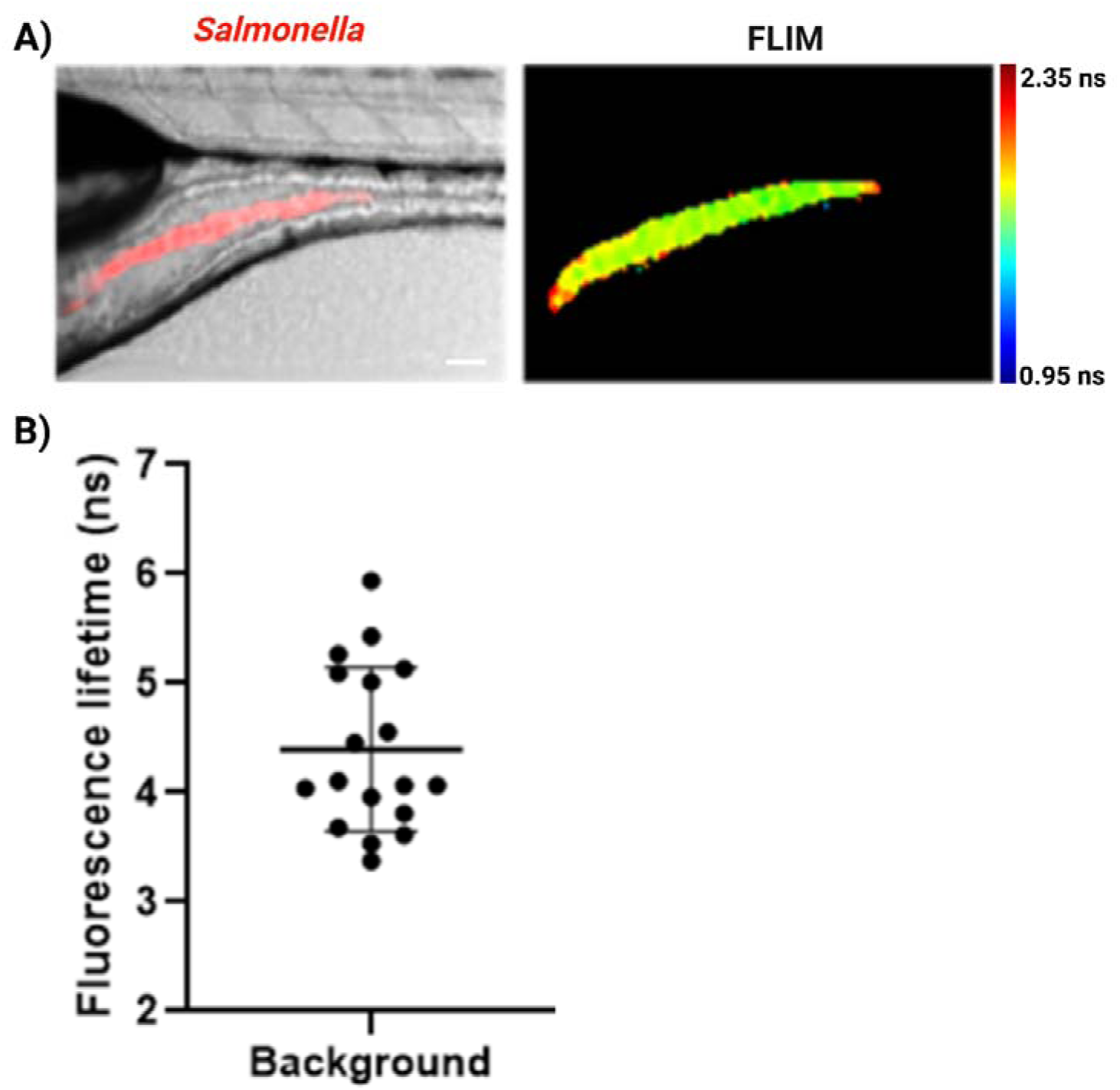
Merged fluorescence microscopy images of the anterior region of zebrafish infected with *Salmonella* with the corresponding FLIM image in the right panel. Scale bars, 50 μm. (B) Quantification of lifetime values for zebrafish background lifetime.

