## Supplementary figures and images for "FLIM Imaging of mCherryTYG Deciphers pH Dynamics and Lifestyles of *Salmonella* Typhimurium"

### Supplementary Movie S1.gif

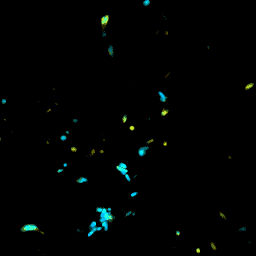
